# Fibrin fragment E potentiates TGF-β-induced myofibroblast activation and recruitment

**DOI:** 10.1101/829945

**Authors:** Peder Fredlund Fuchs, Femke Heindryckx, Carlemi Calitz, François Binet, Sara Marie Øie Solbak, Helena U. Danielson, Johan Kreuger, Pär Gerwins

## Abstract

**Background:** Fibrin is an essential constituent of the coagulation cascade, and the formation of hemostatic fibrin clots is central to wound healing. Fibrin clots are over time degraded into fibrin degradation products as the injured tissue is replaced by granulation tissue.

**Objectives:** study the role of the fibrin degradation product fragment E (FnE) in fibroblast activation and migration.

**Methods:** Rat kidney fibroblasts (NRK-49F), human fetal lung fibroblasts (HFL1) and immortalized human foreskin fibroblasts (BJ) were exposed to FnE and TGF-β. Fibroblast activation was measured via quantitative RT-PCR and western blot for α-SMA and Collagen Iα1. A microfluidic chemotaxis assay was used to study directional migration. To investigate if FnE interacts with integrins, direct binding experiments were performed using surface plasmon resonance biosensor technology. Infusion pumps releasing FnE were implanted subcutaneously in mice and profibrotic effects of FnE were analyzed using immunohistochemical staining of the wound area.

**Results:** We present evidence that FnE is a chemoattractant for fibroblasts and that FnE can potentiate TGF-β-induced myofibroblast formation. FnE forms a stable complex with αVβ_3_ integrin, and the integrin β_3_ subunit is required both for FnE-induced fibroblast migration and for potentiation of TGF-β-induced myofibroblast formation. Finally, subcutaneous infusion of FnE in mice results in a fibrotic response in the hypodermis. These results support a model where FnE released from clots in wounded tissue promote wound healing and fibrosis by both recruitment and activation of fibroblasts. Fibrin fragment E could thus represent a therapeutic target for treatment of pathological fibrosis.

**Essentials:**

– Fibrin is an essential constituent of the coagulation cascade, and fibrin clotting is central to wound healing.
– Fibrin clots are over time degraded into fibrin degradation products, including fibrin fragment E (FnE).
– FnE is a chemoattractant for fibroblasts and can potentiate TGF-β-induced myofibroblast formation.
– FnE released from clots in wounded tissue could promote wound healing and fibrosis.

## Introduction

The coagulation system is involved in a wide range of inflammatory conditions, such as rheumatoid arthritis (1), scleroderma (2), liver cirrhosis (3) and sepsis (4). Initially, tissue injury results in the formation of a cross-linked fibrin matrix that creates hemostasis and provides a provisional matrix for wound repair and tissue regeneration. Platelets trapped in the fibrin mesh release chemotactic factors such as platelet-derived growth factor (PDGF-BB), which together with other growth factors, recruits fibroblasts and inflammatory cells (5, 6). The injured tissue is replaced by granulation tissue, which is a collagen rich matrix with fibroblasts and a dense network of new vessels. Upon activation, fibroblasts become contractile myofibroblasts that express α-smooth muscle actin (α-SMA) and secrete large quantities of extracellular matrix proteins, such as collagen I. Myofibroblasts are mainly, but not solely, derived from nearby resting tissue fibroblasts (7) and have been shown to be necessary for normal wound healing (8). The transition from fibroblast to contractile myofibroblast is mediated by TGF-β, together with extra domain A containing fibronectin and tensile stress. Myofibroblasts play an essential role in different medical conditions such as fibrosis and chronic inflammatory diseases (9, 10). Myofibroblasts are also major constituents of the tumor stroma, where they likely promote disease progression (11).

Coagulation is initiated by the activation of thrombin, which mediates cleavage of circulating fibrinogen to generate fibrin, which polymerizes to form a dense fibrin meshwork (12). To accommodate the expanding granulation tissue and the subsequent scar tissue formation, fibrin is continuously degraded by plasmin, generating fibrin degradation products (FDPs) (13, 14). In the clot, each fibrin monomer contains a central E-domain flanked by one D-domain on each side, and each D-domain is cross-linked to a neighboring D-domain creating a continuous fiber (15). In order to degrade the clot plasmin is generated by cleaving of the precursor plasminogen. Plasmin cleaves fibrin between the E- and D-domain, which generates the FDPs, fibrin fragment E (FnE) and D-dimer (FnDD) (14, 16, 17). Patients with chronic inflammatory diseases often exhibit increased serum levels of FDPs, which has been correlated with disease progression and poor prognosis in, for instance liver cirrhosis (3, 18), systemic scleroderma (19) and pulmonary fibrosis (20). Fibrin degradation products have been shown to have biological effects *in vitro* (21, 22) as shown by experiments using endothelial cells (22, 23), macrophages (24), vascular smooth muscle cells (21, 22, 25) and leukocytes where they increase cell migration (21, 23, 25), inflammatory response (24), proliferation and differentiation (23) and regulate survival and apoptosis (23). However, the effects of FnE on fibroblast migration and differentiation into myofibroblasts have not been investigated previously.

In this study we show that FnE is a chemotactic factor for fibroblasts and that FnE potentiates TGF-β-induced myofibroblast activation and induces a profibrotic response when infused in the skin of mice. These results show the importance of the coagulation cascade in wound healing and fibrosis, which could be therapeutically relevant for patients with pathologic inflammatory conditions.

## Results

### FnE potentiates expression of α-SMA induced by TGF-β

To study if degradation products released from fibrin gels could affect α-SMA expression in fibroblasts, a transwell assay was employed with fibroblasts embedded in a fibrin gel in the insert and effects on fibroblasts at the bottom of the well examined (Figure 1A). First, TGF-β was shown to induce α-SMA expression in human fetal lung fibroblasts (HFL1 and the expression of α-SMA could be decreased by the addition of serine protease inhibitor aprotinin (Figure 1B, C). This experiment suggested that FDPs released from the gel by cell-derived proteases stimulated TGF-β –induced α-SMA expression in an additive or synergistic manner. This possibility was further investigated using purified FnE, which when added to HFL1 cells potentiated TGF-β-induced α-SMA expression by around two-fold in a dose-dependent manner (Figure 1D, E, F). Notably, FnE did not alter the basal α-SMA expression levels when added alone (Figure 1D). The effect of FnE reached a maximum at 2 nM after 48 h (Figure 1H). Addition of a fixed concentration of FnE (2 nM) together with increasing concentrations of TGF-β revealed that the synergistic effect was present at 5 ng/ml TGF-β, but not at lower concentrations (Figure 1G). FnE was also shown to potentiate TGF-β-induced α-SMA expression in immortalized human foreskin fibroblasts (BJ hTERT, Figure 1H) and rat kidney fibroblasts (NRK-49F, Figure 1I, J), suggesting that the response to FnE is a basic feature of fibroblasts.

**Figure 1.**
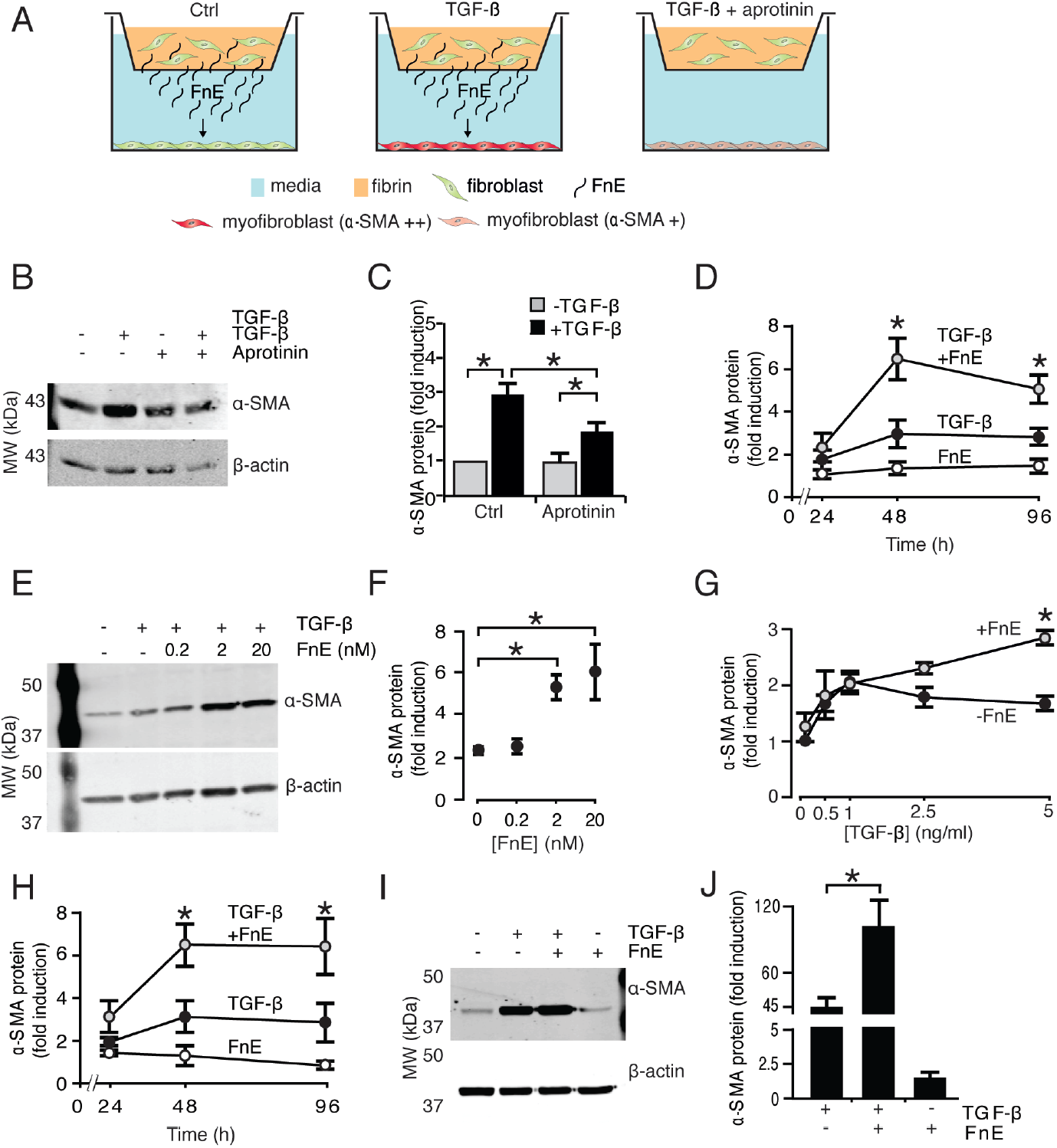
FnE synergistically enhance TGF-β-induced α-SMA protein expression in a time- and dose-dependent manner. (A) Human HFL1 fibroblasts were grown in transwell inserts in a fibrin gel, as well as on the bottom of the well, and treated as indicated. (B) Representative western blot for α-SMA obtained from cells growing on the bottom of the well 48 h after stimulation with or without TGF-β (5 ng/ml). (C) Quantification of the results in B normalized to β-actin and expressed as fold increase compared to unstimulated control. (D) HFL1 fibroblasts were starved and cultured in the presence of either TGF-β (5 ng/ml), or FnE (2 nM), or both factors for the indicated periods of time. α-SMA levels were determined by western blotting and normalized to β-actin and expressed as fold increase compared to control cells. (E) Representative western blot showing the dose-dependent potentiation of TGF-β-induced α-SMA expression by FnE and (F) quantification of the results normalized to β-actin and expressed as fold increase compared to controls. (G) α-SMA protein levels in HFL1 cells cultured with or without FnE (2 nM) and increasing concentrations of TGF-β. (H) α-SMA protein levels in human BJ hTERT fibroblasts treated either with TGF-β or with FnE, or a combination of both. (I) Representative western blot showing α-SMA protein expression in rat NRK-49F fibroblasts cells after a 48 h stimulation with TGF-β (5 ng/ml), FnE (2 nM) or a combination of both and (J) quantification of the results normalized β-actin and expressed as fold increase compared to controls. * p<0.05. Error bars represent standard error of the mean (*n* ≥ 3).

In order to elucidate if other FDPs than FnE could impact α-SMA expression, we investigated the effects of adding the E fragment from non-cross-linked fibrinogen (FgnE) or D-dimer (FnDD) from cross-linked fibrin to HFL1 cells. At equimolar concentrations, FgnE, had no effect while FnDD potentiated TGF-β induced α-SMA expression, although to a lesser extent than FnE (Figure 2A, B). Neither FnE nor FgnE nor FnDD altered the basal α-SMA expression levels. Further, FnDD could not enhance α-SMA expression when added together with FnE (Figure 2C).

**Figure 2.**
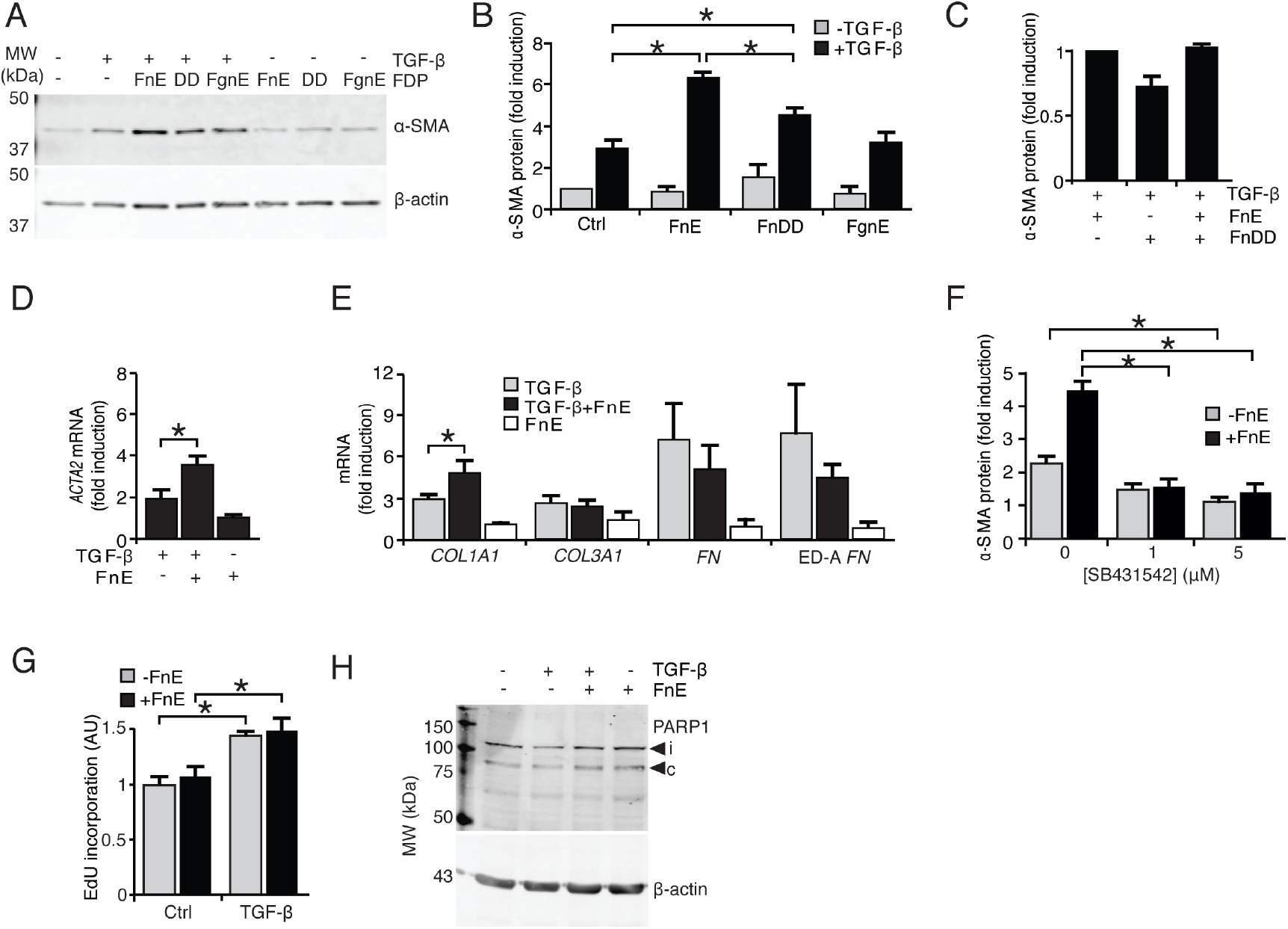
FnE and FnDD but not FgnE enhance TGF-β-induced myofibroblast activation. (A) Representative western blot and (B) quantification of α-SMA protein levels in HFL1 cells cultured for 48 h with TGF-β (5 ng/ml) alone or in combination with different FDPs (2 nM). Values in (B) are normalized to β-actin and expressed as fold increase compared to untreated control cells cultured in starvation medium. (C) α-SMA protein expression in HFL1 cells after a 48 h stimulation with TGF-β in combination with FnE and/or FnDD. (D) Relative α-SMA (*ACTA2*) mRNA expression and (E) other major matrix component transcripts in HFL1 cells after a 48 h treatment with the indicated factors (TGF-β, 5 ng/ml; FnE, 2 nM); collagen Iα1 (*COL1A1*), collagen IIIα1 (*COL3A1*), fibronectin (*FN*) and extra domain A containing fibronectin (EDA *FN*). (F) TGF-β (5 ng/ml) and TGF-β + FnE (2 nM) induced α-SMA protein expression in HFL1 cells is inhibited by a 48h treatment using the TGF-β receptor kinase inhibitor SB431542. (G) Proliferation in response to TGF-β (5 ng/ml), FnE (2 nM) or a combination of both for 48 h, expressed as the fraction of EdU positive nuclei in treated cells compared to unstimulated control cells. (H) Western blot showing the levels of PARP-1 in HFL1 cells treated or not with TGF-β and FnE; molecular species corresponding to the cleaved (c) and intact (i) forms are indicated by arrows. *p<0.05. Error bars represent standard error of the mean (*n* ≥ 3).

Collagen I, collagen III, fibronectin and extra domain A fibronectin (ED-A FN) are, in addition to α-SMA, proteins that are associated with myofibroblast differentiation. Besides α-SMA mRNA (*ACTA2*), FnE also increased TGF-β-stimulated Collagen Iα1 mRNA (*COL1A1*) but not Collagen III α1 (*COL3A1*), fibronectin (*FN*) or ED-A containing fibronectin (*ED-A FN*) in HFL1 cells (Figure 2 D, E). Inhibition of TGF-β receptor signaling by the low molecular inhibitor SB431542 blocked both TGF-β-induced as well as FnE-mediated potentiation of α-SMA expression in HFL1 cells (Figure 2F), demonstrating that TGF-β signaling is critical in order for FnE to exert its effect on α-SMA potentiation.

### FnE does not influence cell proliferation or induce apoptosis

It has been reported that FnE stimulates endothelial cell proliferation (23) and induces apoptosis (26). Here, TGF-β caused a slight increase in proliferation of HFL1 cells but FnE alone, or in combination with TGF-β, had no effect (Figure 2G). Treatment of HFL1 cells with FnE did not increase apoptosis as measured by increased cleavage of the DNA-repair enzyme PARP-1, which is commonly used as an indication of apoptosis (27) (Figure 2H).

### FnE is chemotactic for fibroblasts

FnE has previously been reported to induce migration of endothelial and vascular smooth muscle cells. We therefore investigated the chemotactic effects of FnE on fibroblasts. A microfluidic cell migration chamber (28, 29) was used to analyze the effects of stable gradients of FDPs on chemotaxis and chemokinesis of individual cells using time-lapse microscopy. FnE was the only FDP that had a chemotactic effect (Figure 3A, B) similar to that of the positive control PDGF-BB. The net migration distance (irrespective of direction) was similar for all FDPs investigated (Figure 3C, D) indicating that none of them induced a chemokinetic response.

**Figure 3.**
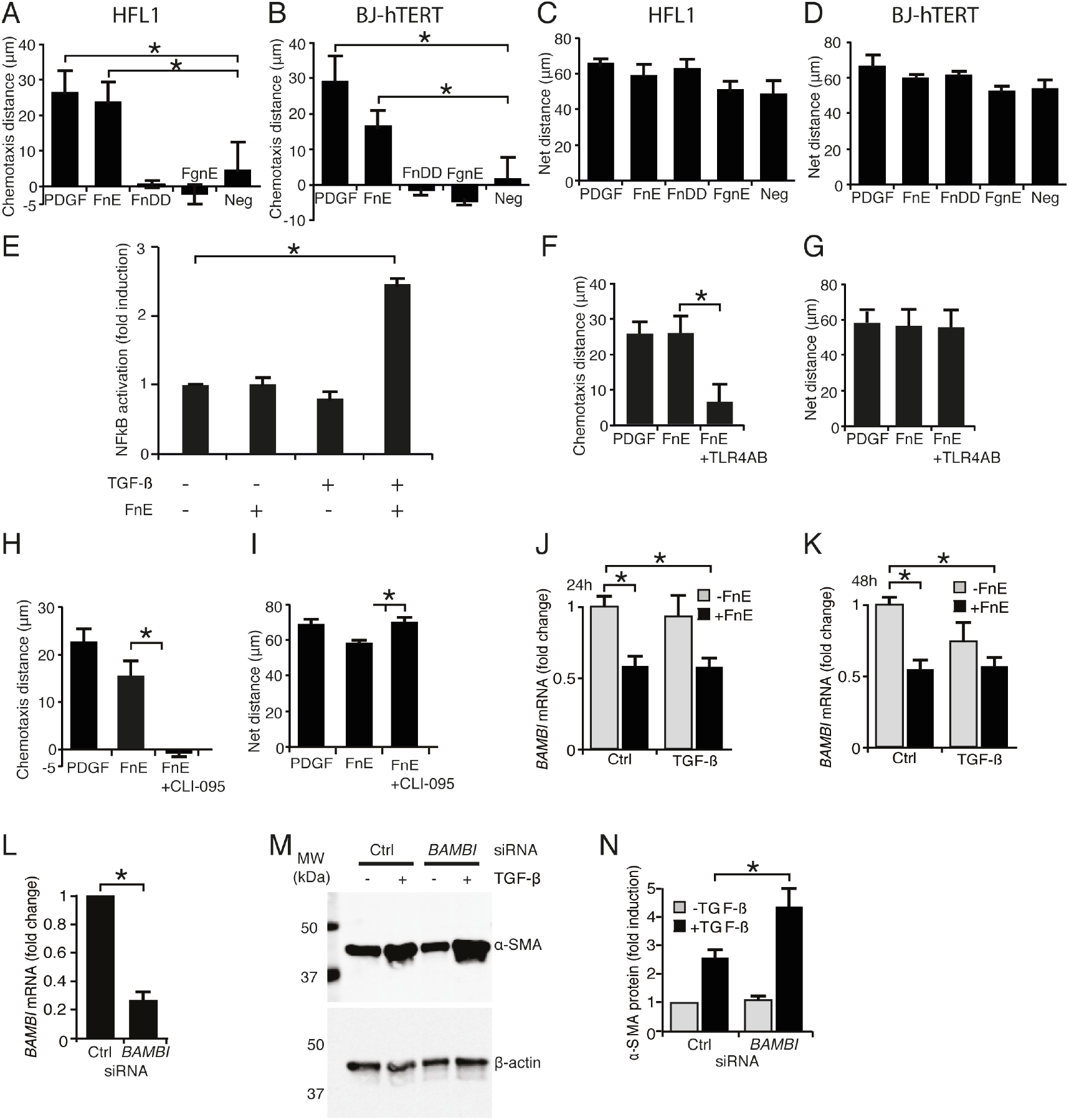
FnE-induced fibroblast chemotaxis is dependent on TLR4 and on interactions with αVβ_3_ integrins. (A) HFL-1 and (B) BJ-hTERT fibroblasts were exposed to stabile gradients of either PDGF-BB (positive control) or the different FDPs using a microfluidic chemotaxis assay to study effects on cell migration. Of the FDPs, only a gradient of FnE elicited a chemotactic response. The net distance migrated by (C) HFL-1 and (D) BJ-hTERT fibroblasts irrespective of directionality was also scored. (E) NFkB activation in HFL1 cells 48 h after stimulation with TGF-β (5 ng/ml), FnE (2 nM) or a combination of both. (F) Chemotaxis and (G) net distance migrated by HFL1 cells in the presence of the TLR4-blocking antibody. (H) Chemotaxis and (I) net distance migrated by HFL1 cells in the presence of the TLR4 inhibitor CLI-095. *BAMBI* mRNA levels in HFL1 cells 24 h (J) and 48 h (K) after treatment with TGF-β (5 ng/ml) and/or FnE (2 nM). (L) Supression of *BAMBI* mRNA levels using siRNA. (M, N) Representative western blot and corresponding quantifications of the α-SMA protein levels in siRNA-treated cells with and without TGF-β (5 ng/ml) stimulation. * = p<0.05. Error bars represent standard error of the mean (*n* ≥ 3).

### FnE-mediated chemotaxis is dependent on Toll like receptor 4 and BAMBI

Toll-like receptor 4 (TLR4) has been reported to bind fibrinogen (30) and thereby to regulate proliferation of fibroblasts (31), which makes TLR4 a candidate receptor for FnE. The combination of TGF-β and FnE lead to a significant activation of NFkB, which could mean that TLR4 is involved in FnE’s potentiation of TGF-β (Figure 3E). Treatment of HFL1 fibroblasts with a specific blocking antibody against TLR4 (Figure 3F, G) and the TLR4 inhibitor CLI-095 (also known as TAK-242) (Figure 3H, I) (32) inhibited the chemotactic effect of FnE (Figure 3H, I). This supported our hypothesis that TLR4 is a candidate receptor for FnE. Bacterial lipopolysaccharide (LPS) has been shown to enhance TGF-β signaling by TLR4 mediated down-regulation of the TGF-β pseudo-receptor BAMBI (33). In agreement, FnE treatment of HFL1 cells caused a 50% reduction in *BAMBI* mRNA levels already after 24h (Figure 3J), which persisted for at least 48h (Figure 3K). Reduction of BAMBI levels with siRNA (Figure 3L) mimicked the effect of FnE with potentiation of TGF-β induced α-SMA expression without addition of FnE (Figure 3M, N).

### FnE-mediated chemotaxis and α-SMA induction depend on β3 integrins

Both fibrinogen and fibrin contain several tripeptide arginine-glycine-aspartate (RGD) sequences, making them potential binding partners for RGD-binding integrins. In order to investigate if FnE interacts with integrins with a biologically relevant mechanism and affinity, direct binding experiments were performed using surface plasmon resonance (SPR) biosensor technology. Thus, FnE was immobilized onto a sensor chip and soluble recombinant human αVβ_3_-integrin injected as an analyte over the surface of the chip (20). αVβ_3_-integrin displayed a concentration-dependent interaction with FnE, forming a stable complex (Figure 4A). This was not observed for FnDD (Figure 4B). The binding curves were not well described by a simple 1:1 binding model, suggesting that the interaction was more complex (Figure 4C). The slow dissociation rates indicate a strong interaction with a dissociation constant (K_D_) ranging in nM scale (2-20 nM dependent on interaction model). To further investigate the potential importance of the RGD element in FnE a cyclic-RGD peptide EMD 66203 (cRGD) was used, which competes for the binding to RGD binding integrins. cRGD treatment eliminated the synergistic effect of FnE on TGF-β-induced α-SMA expression in HFL1 cells, but did not affect the response to TGF-β alone (Figure 4D, E). In contrast, treatment with cRGD increased both chemotaxis as well as the total distance migrated of HFL1 cells in response to PDGF-BB and FnE gradients (Figure 4F, G). Notably, however, the ratio between chemotaxis and the total distance migrated remained similar compared to previous results, indicated by dashed grey lines.

**Figure 4.**
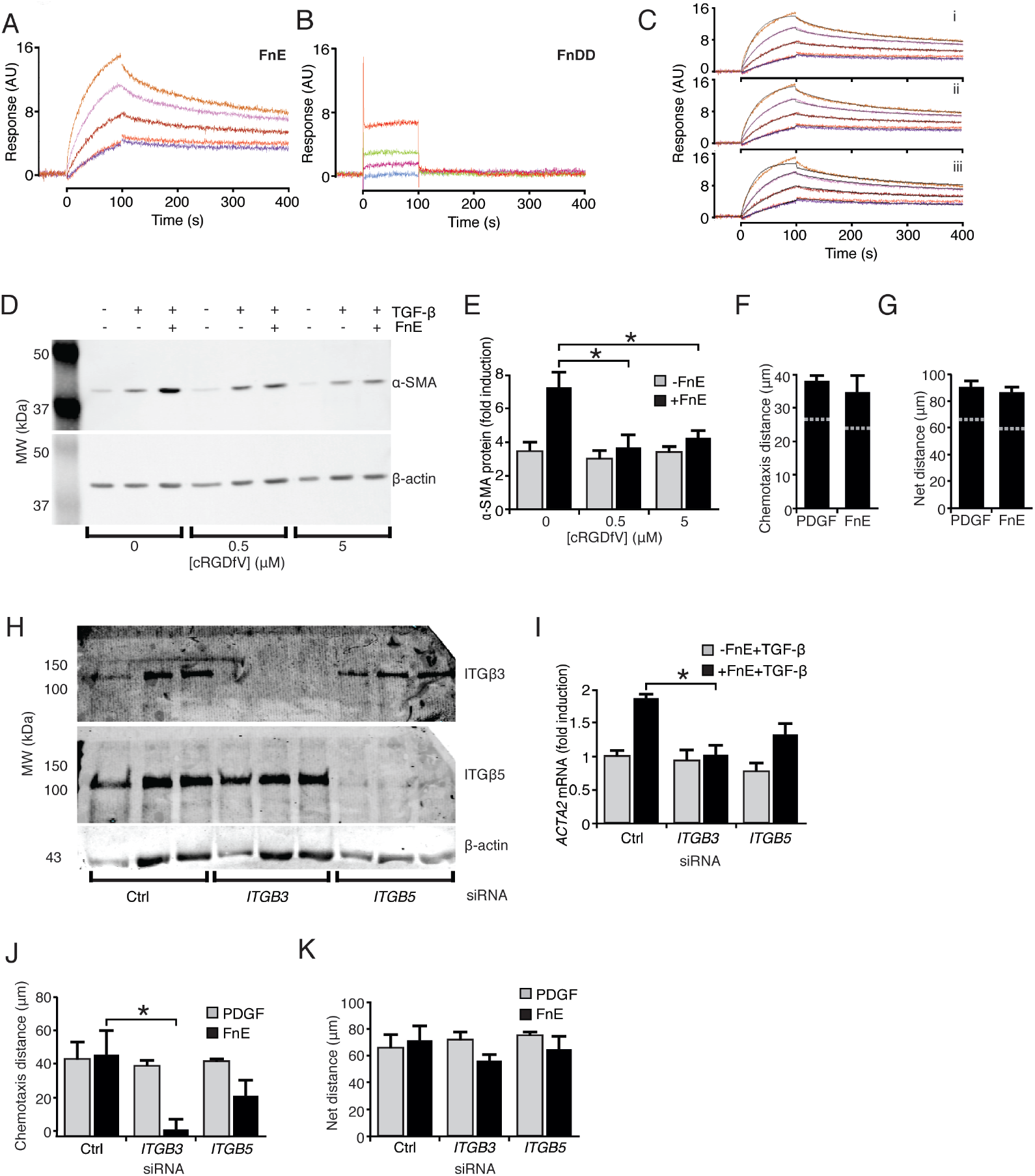
FnE-mediated potentiation of α-SMA expression is dependent on interactions with αVβ_3_ integrins. Interactions between soluble recombinant αVβ_3_ (5-50 nM) and immobilized FnE (A) or FnDD (B) were determined by SPR. Immobilization was at 952 RU and 1251 RU respectively. (C) SPR sensorgrams of the interaction between FnE and integrin αVβ_3_ fitted to a bivalent analyte model (i), a heterogeneous ligand model (ii) or to the 1:1 two state interaction model (iii). (D) Representative western blot and (E) quantification of α-SMA protein levels of HFL1 fibroblasts in the presence of the cyclic-RGD peptide EMD 66203 in combination with TGF-β (5 ng/ml) and/or FnE (2 nM). α-SMA was normalized against β-actin and expressed as compared to vehicle (DMSO) treated controls. (F) Chemotaxis and (G) net migration of HFL1 cells in presence of 5 µM cyclic-RGD peptide. Dashed grey lines indicate mean values for cells without addition of cyclic-RGD. (H) Integrin β3 and β5 protein levels in HFL1 cells after transfection with siRNA selectively targeting these integrins, as indicated. (I) α-SMA (*ACTA2*) mRNA levels of siRNA transfected cells stimulated with TGF-β (5 ng/ml) with or without addition of FnE (2 nM) for 48 h. (J) Chemotaxis and (K) net migration of HFL1 cells treated with siRNA to suppress integrin β3 or β5 in response to a PDGF-BB (0-20 ng/ml) or FnE (0-2 nM) gradients. * p<0.05. Error bars represent standard error of the mean (*n* ≥3).

The cRGD peptide targets the αVβ_3_–integrin complex as well as the αVβ_5_-integrin complex (34). In order to elucidate if both of these complexes are involved in mediating the effects of FnE we performed siRNA knockdown experiments targeting either integrin β3 or β5 in HFL1 cells. Robust reductions in integrin β3 and β5 protein levels as a result of siRNA treatment were confirmed by western blotting (Figure 4H). siRNA-mediated suppression of integrin β3, but not β5, abolished the potentiating effect of FnE on TGF-β-induced α-SMA mRNA expression (Figure 4I). A reduction of the integrin β3 levels also diminished the chemotaxis toward FnE whereas a reduction of the integrin β5 levels had no significant effect on FnE-induced chemotaxis (Figure 4J). Notably, PDGF-BB-induced chemotaxis as well as general cell motility in any condition was not affected by suppression of β3 or β5 integrins (Figure 4J, K).

### Profibrotic effects of FnE in vivo

In order to validate these *in vitro* results, we used an *in vivo* mouse model where profibrotic effects are analyzed after continuous subcutaneous infusion of FnE by mini-osmotic pumps. A single pump loaded with FnE or saline (control) was implanted subcutaneously on the back of C57/BL6 mice and kept in place for 2 weeks. Immunohistochemical analysis of sections of skin taken from the area in proximity to the infusion site revealed that FnE-treated mice displayed an increased number of α-SMA-positive myofibroblasts in the hypodermis (Figure 5A, B, C).

**Figure 5.**
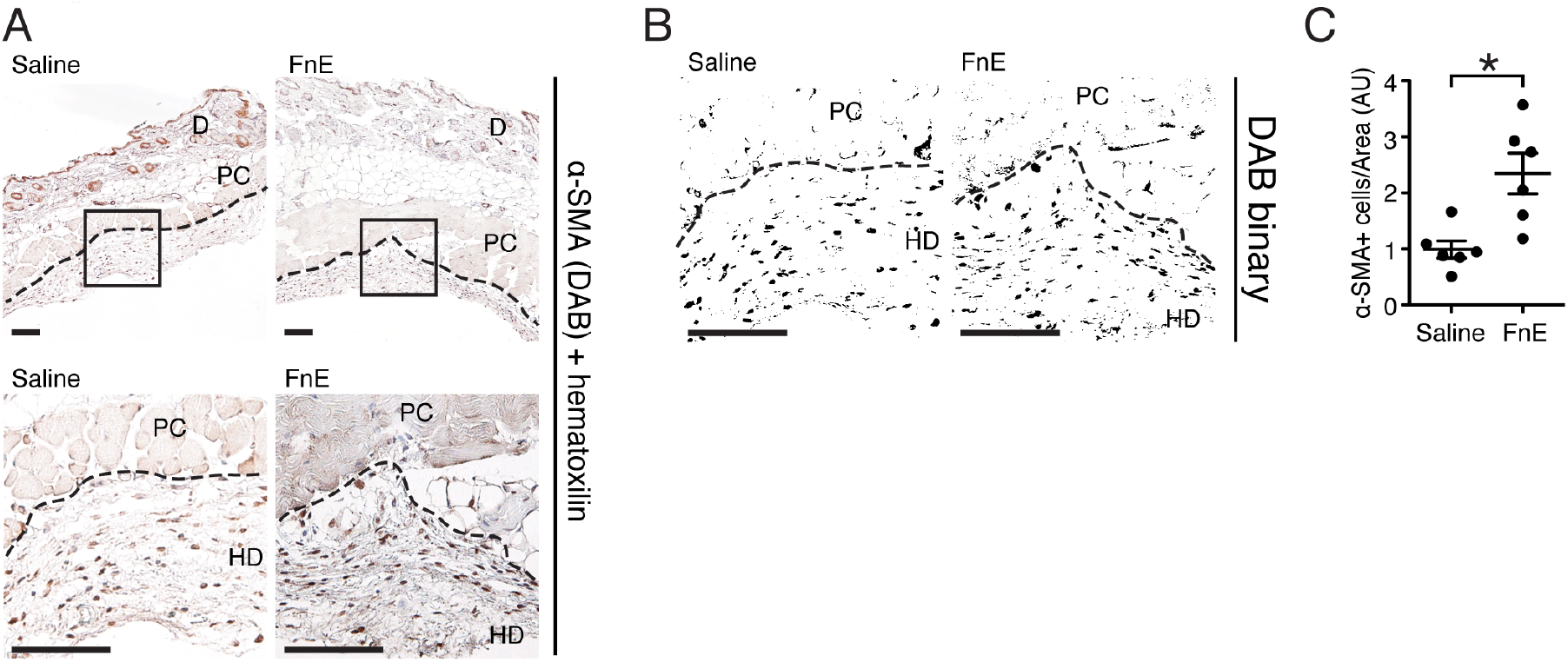
FnE-infusion results in increased numbers of α-SMA positive cells in the hypodermis of mice *in vivo*. Mini-osmotic pumps containing either saline (control) or FnE (20 µg) was implanted subcutaneously in C57Bl/6 mice and left in place for 2 weeks. (A) α-SMA-positive cells in mouse hypodermis (HD) were identified by immunohistological staining near the infusion site. D, dermis; PC, panniculus carnosus; Color de-convolution and conversion to binary images (B) was performed to enable quantification of α-SMA-positive cells. (C) α-SMA+ cells/area HD was quantified and normalized to the mean value in the saline control group. Scale bar 100 µm, * p<0.05. Individual mice are represented by dots. Error bars represent standard error of the mean (*n* = 6 per group).

## Discussion

Deposition of a fibrin-rich matrix in the interstitium is an integral part of tissue repair and fibrin deposits are in addition found in association with tumors (11, 35), rheumatoid arthritis (36), inflammatory diseases and fibrotic conditions (3, 18–20, 37) where fibrin has been proposed to promote disease development. During tissue injury, circulating fibrinogen will generate a dense network of cross-linked fibrin upon activation of the coagulation cascade. This network provides a scaffold that directly interacts with cells to create a hemostatic plug. The fibrin scaffold also provides a provisional matrix for wound repair and tissue regeneration. The continuous degradation of fibrin generates the FDPs, fibrin fragment E and D-dimer. In this study we provide evidence for that low nanomolar concentrations of FnE stimulates fibroblast migration comparable to that of PDGF-BB which is a strong chemoattractant (38), and potentiates TGF-β induced myofibroblast formation.

FnE, in contrast to polymerized fibrin, is freely diffusible and we suggest that FnE released from polymerized fibrin by proteolysis forms a concentration gradient that recruits fibroblasts to the injured tissue as demonstrated by our live-cell tracking experiments. As the fibroblast move closer to the fibrin clot they become increasingly activated by higher levels of FnE and other growth factors and cytokines. The initial recruitment of fibroblasts is likely also dependent on local and acute release of PDGF-BB from platelets, while FnE may act to continuously recruit fibroblasts as long as there is degradation of fibrin deposits.

One factor in particular that fibroblasts encounter as they migrate into the wounded area is TGF-β. This potent cytokine is produced by neutrophils and macrophages present in the wound, as well as by keratinocytes and fibroblasts. As demonstrated here, FnE will be capable of enhancing TGF-β-induced α-SMA expression and collagen I production. Notably, as FnE does not induce the transition towards a myofibroblast phenotype by itself, only fibroblasts that are migrating into TGF-β-rich areas of the wound would become fully differentiated into myofibroblasts. Accordingly, when mice were implanted with mini-osmotic pumps that continuously released FnE, a significant increase of α-SMA positive cells was observed near the implantation site, which indeed is a wounded area due to the surgical procedure that was needed to implant the pumps.

Interestingly, FnE down-regulated BAMBI, which is a pseudoreceptor that interacts with TGF-β and BMP type II receptors (39) to form a non-signaling receptor complex that negatively modulates TGF-β responses. Reduced BAMBI expression potentiated TGF-β-induced responses. These results are in line with a previous study where TGF-β-induced α-SMA and collagen I expression was enhanced by down-regulation of BAMBI (40).

The Toll-like receptors have until recently been considered to mediate only innate immune responses but have lately been shown to bind endogenous ligands such as hyaluronan (41), HMGB1 (42) and fibrinogen (30). Although TLR4 is mainly expressed by immune cells as part of the innate immune system, TLR4 activation has also been shown to occur in non-immune cells within injured tissue micro-environments, such as renal fibrosis (43) and scleroderma (44, 45). Our data now suggests that TLR4 modulates FnE-stimulated fibroblast migration. Fibroblast chemotaxis in response to FnE also requires integrin β_3_ as determined here by siRNA knockdown experiments. αVβ_3_ integrin was also found to bind directly to FnE, forming a very stable complex (Fig. 4). In agreement with this, the synergistic effect of FnE on TGF-β induced α-SMA was dependent on integrin αVβ_3_ as shown by using the cRGDfV peptide and by knockdown of β_3_ with siRNA. The finding that inhibition of αVβ_3_ and αVβ_5_ using cRGDfV did not reduce the FnE induced chemotaxis may be due to that there, at any given time, may be enough free αVβ_3_ to distinguish between the high and low concentration end of the FnE gradient.

Subcutaneous infusion of FnE into mice induced a profibrotic response with increased number of α-SMA positive myofibroblasts in the hypodermis. These results suggest a role for FnE in fibrotic conditions in agreement with the proposed functions of fibrin, fibrinogen and FDPs in wound healing and inflammation. In wound healing, FnE might promote recruitment and differentiation of myofibroblasts and enhance the formation of granulation tissue. FnE could also contribute to revascularization through looping angiogenesis where myofibroblast mediated contraction of the provisional matrix creates biomechanical forces that mediate and direct neovascularization (46). In addition, fibrin antibodies have also been shown to directly inhibit angiogenesis, thus further indicating a role for FnE in neovascularization (47), which is an essential aspect in wound healing, but also contributes to progression and metastasis of solid tumors. The release of FnE from fibrin deposits in wounds and tumors would form stable gradients to direct recruitment of fibroblasts as long as fibrin deposits are present. Tumors have historically been called “a wound that does not heal” (48). Given the deposition of fibrin and the disruption of coagulation pathways found within many tumors, there is possible that FnE may be involved in tumor progression by recruiting cancer-associated fibroblasts (49). These fibroblasts are known to promote tumorigenesis, by initiating the remodeling of the extracellular matrix and by secreting cytokines (11). In support, there are indications that cancer patients treated with anti-coagulants have an increased survival, which is not only due to a decreased incidence of thromboembolic events (50–52). The findings in animal models that an antibody against FnE inhibits tumor growth (53) and that metastasis is reduced in fibrinogen deficient animals (54) further supports such a concept.

Accumulating evidence suggests a role for fibrinogen, fibrin and FDPs in fibrotic disease. Fibrinogen-deficient mice are protected from kidney fibrosis in a model of unilateral urethral obstruction with reduced number of fibroblasts, lower levels of α-SMA expression and less collagen I production (31). Further evidence for a role in fibrosis is provided by the finding that injection of fibrin into the tunica albuginea in mice induces fibrotic response similar to Peyronie’s disease in humans (55). Fibrin deposition is also one of the most conspicuous and consistent features of both human rheumathoid arthritis and experimental animal models of arthritic disease (56).

The current study shows that FnE at low nanomolar concentrations is a potent inducer of fibroblast chemotaxis, enhances myofibroblast differentiation, and is profibrotic *in vivo*. These results further expand the role of fibrin beyond hemostasis, and may have implications for the understanding of the pathophysiology of wound healing, fibrosis and tumor growth.

## Materials and methods

### Cell culture and reagents

Rat kidney fibroblasts (NRK-49F, ECACC), human fetal lung fibroblasts (HFL1, ECACC) and immortalized human foreskin fibroblasts (BJ hTERT) were routinely cultured in MEM supplemented with non-essential amino acids, 25 mM HEPES (Sigma), 10 % FBS (Gibco), 2 mM glutamax (Gibco) and 1 mM sodium pyruvate (Gibco). Cells were cultured at 37°C in a humidified atmosphere containing 5% CO_2_. Growth medium without FBS was used for starvation, stimulation and cell migration experiments.

For experiments where cells were stimulated, cells were first detached using trypsin-EDTA (Gibco), thereafter resuspended in growth medium and plated at a density of 5×10^3^ cells/cm^2^. Cells were allowed to attach for 24 h before being starved for 24 h where after fresh starvation medium containing TGF-β and/or FDPs and inhibitors were added. Cells were harvested at indicated times for RNA or protein extraction. RNA was isolated using Total RNA Kit I (EZNA). For protein extraction, the cells were lysed either in protein lysis buffer (150 mM NaCl, 20 mM Hepes, 10 % glycerol and 1% NP-40) or directly in SDS-containing sample buffer (see SDS-PAGE and western blot), both solutions containing 100 U/ml aprotinin, 1 mM Pefablock, 2.5 mM EDTA and 100 nM sodium vanadate was added. TGF-β was added at 5ng/ml, and FDPs at 2 nM unless otherwise indicated. The inhibitors, SB431542 and the cRGDfV peptide EMD 66203 (both from Sigma-Aldrich) were added 30 minutes prior to stimulation.

### Fibrin gel degradation

HFL1 cells were detached by trypsin-EDTA, resuspended in serum free MEM and pelleted by centrifugation to remove the trypsin. The pellet was resuspended in serum free MEM containing 2.5 mg/ml bovine fibrinogen (Sigma) and, for some conditions, 50 U/ml aprotinin. Recombinant human thrombin (1 U/ml, Sigma) was added to the bottom of a 3.0 µm costar polycarbonate transwell membrane insert (Corning) and HFL-1 cells suspended in 500 µl fibrinogen solution was added to the insert, mixed, and allowed to polymerize for 1 h at 37°C. After polymerization, serum-free MEM was added on top of the gel. The inserts were after 6 h transferred to new wells containing pre-starved HFL1 cells and left overnight before indicated factors was added to the lower well. Aprotinin was added to maintain final concentration of 50 U/ml. Cells in the lower well were harvested 48 h post stimulation.

### Electroporation of cells for siRNA transfection

HFL1 cells were detached using trypsin-EDTA solution (Gibco) and approx. 600 000 cells pelleted and resuspended in 100 µl transfection solution (Mirus Bio) containing 400 pg targeting siRNA or non targeting control siRNA. The cell suspension was transferred to an electroporation cuvette (Mirus Bio) and electroporation was carried out using the Nucleofector I (Amaxa) program T-23 or S-05. After electroporation, the cells were divided and seeded equally into 3 wells in a 6-well plate. Cells were allowed to settle for 6-8 h before being starved over night in serum-free medium. Stimulation with the indicated factors was initiated 48 h before harvest (at 72 h post transfection). Silencer select siRNAs (Ambion) used were targeting either *ITGB3* (integrin β3, siRNA id s7580 and s7582), *ITGB5* (integrin β5, siRNA id s7590 and s7589) and *BAMBI* (siRNA id s24523). Negative control siRNA #1 (AM4635) was used as non-targeting control.

### SDS-PAGE and western blot

Protein lysates in lysis buffer were mixed with 2x sample buffer (0.1 M Tris, 10% glycerol, 2% SDS, 5% β-mercaptoethanol and 50 µg/ml bromophenol blue) and heated to 95°C for 5 minutes before being loaded onto a 10% polyacrylamide gel. After separation, the proteins were transferred to an Immobilon-Fl membrane (Millipore). The membrane was after transfer blocked using the Odyssey blocking buffer (Licor) diluted 1:4 in PBS, and then incubated sequentially with primary and secondary antibodies. After primary and secondary antibody incubation the membrane was washed 3×15 minutes in PBS-T (Phosphate buffered saline (Gibco), 0.1% Tween-20). Primary antibodies used were mouse anti-α-SMA (clone 1A4, Sigma-Aldrich) 1:10 000, rabbit anti-β-actin (ab8227, Abcam) 1:5000, mouse anti-PARP1 (BD-bioscience) 1:1500, mouse anti-integrin β3 (Cell signaling) 1:500, rabbit anti-integrin β5 (Abnova) 1:250, all added in blocking buffer with 0.1% Tween-20. Secondary antibodies used were: goat-anti-mouse alexa 680 (Invitrogen) and goat-anti-rabbit IRDye 800 (Rockland) 1:20 000 diluted in blocking buffer with 0.1% Tween-20 and 0.01% SDS. All incubations were carried out at room temperature for 1 h or over night at 4°C. The membranes were scanned using an Odyssey scanner (LI-COR Biotechnology) and band intensities quantified using the Odyssey 2.1 software and normalized to the β-actin signal in each sample.

### RT PCR and qPCR

Total RNA was extracted using the Total RNA Kit I (EZNA) and cDNA synthesis was carried out using iScript (Biorad) according to the manufacturers instructions and using 300 ng total RNA for each sample.

Real-time quantitative PCR (qPCR) was performed using Ssofast Evagreen (Biorad) on a mini Opticon machine (Biorad). Data was analyzed using Bio-rad CFX manager software using the ddCt method and normalized to beta actin. The primer sequences used were hsa *ACTB* (β-actin) 5’-forward TCTACCAATGAGCTGCGTGTG-3’, hsa *ACTB* reverse 5’-AGCCTGGATAGCAACGTACA-3’, hsa *ACTA2* (α-smooth muscle actin) forward 5’-TTCAATGTCCCAGCCATGTA-3’, hsa *ACTA2* reverse 5’-GCAAGGCATACCCTCATAG-3’, hsa *COL1A1* (collagen IαI) forward 5’-GATGGACAGCCTGGTGCT-3’, hsa *COL1A1* reverse 5’-ACCAGGTTCACCGCTGTTAC-3’, hsa *COL3A1* (collagen IIIαI) forward 5’—3’, hsa *COL3A1* reverse 5’—3’, hsa *FN* (fibronectin) forward 5’-TTTGTGGTCTCCTGGGTCTC-3’, hsa *FN* reverse 5’-AGAAGTGGCTGTGCTTGGAA-3’, ED-A containing hsa *FN* forward 5’-AATCCAAGCGGAGAGAGTCA-3’, ED-A containing hsa *FN* reverse 5’-GGTCACCCTGTACCTGGAAA-3’, hsa *ITGB3* (integrin β3) forward 5’-ATGGGGACACCTGTGAGAAC-3’, hsa *ITGB3* reverse 5’-ACGCTTGCAGGTATTTTCG-3’, hsa *ITGB5* (integrin β5) forward 5’-GCCTTGCTTGGAGAGAAAT-3’, hsa *ITGB5* reverse 5’-AATCTCCACCGTTGTTCCAG-3’. hsa *BAMBI* forward 5’-ATCGCCACTCCAGCTACATC-3’, hsa *BAMBI* reverse 5’-GGCAGCATCACAGTAGCATC −3’.

### EdU proliferation assay

The proliferation assay was carried out in medium containing 2% FBS. Cells were incubated with 1 µM 5-ethynyl-2’-deoxyuridine (EdU, Invitrogen) for 6 h (42-48 h post stimulation) before fixation in 4% PFA in PBS for 15 minutes at room temperature. EdU incorporation was visualized using the Click-iT™ EdU Alexa Fluor® 488 or 555 Imaging Kit (Invitrogen), using Hoechst 33342 as nuclear staining, according to the manufacturer’s instructions. Images were obtained using an Axiovert 200M microscope (Zeiss) and analyzed with ImageJ software in order to determine the fraction of EdU positive nuclei. Images of 6 random fields were analyzed per treatment and experiment.

### Microfluidic chemotaxis assay

Migration in response to a stabile gradient of factors was assessed as previously described (28, 29). In short, cells were seeded onto a gelatin coated 35 mm tissue culture dish (BD Falcon), allowed to attach and starved over night after which the microfluidic chamber was applied on top and a gradient generated using indicated factors in serum free medium. Factors gradients used were PDGF-BB (Peprotech) 0-20 ng/ml, FnE, FgnE and FnDD (Hyphen biomed) 0-2 nM over a distance of 500 µm with a flow rate of 1.5 µl/minute. Cell movement was recorded using an Axiovision 200M microscope (Zeiss) for 4 h and tracked using Axiovision software (Zeiss). Cells were kept at 37°C with 5% CO_2_ in a humid atmosphere during the experiments. When inhibitors were used, cells were pre-incubated 1 h before assembly of the migration chamber, followed by presence of the inhibitor in the entire chamber for the entire duration of the experiment. For siRNA treated cells the assay was initiated 46-50 h post nucleofection. Data is presented as chemotaxis distance, measuring movement towards the high end of the gradient, and net distance, measuring the average distance of migration between start- and end points of the individual cells.

### SPR biosensor analyses

SPR biosensor measurements were performed at 25 °C using a Biacore S51 instrument (Biacore AB, Uppsala, Sweden). FnE and FnDD were immobilized with standard amine coupling procedure to a CM5 chip using sodium acetate pH 4 as immobilization buffer to approximately 950 response units (RU) and 1251 (or 5226 RU), respectively. PBS buffer pH 7.4 with 0.005% P20 was used as running buffer, and integrin αVβ_3_ was run as analyte in four different concentrations using a flow rate of 30 µl/min. The surface was regenerated with glycine-HCl pH 2 between each cycle for the interaction with FnE and integrin αVβ_3_. The sensorgrams were corrected for buffer bulk effects and unspecific binding of the samples to the chip matrix by blank and reference surface subtraction (subtraction of inactivated and deactivated flow cell channel). The data were analyzed using Biacore T200 evaluation software version 1.0. The sensorgrams of the interaction between FnE and integrin αVβ_3_ were fitted to several binding models, including 1:1 (Langmuir) binding model, two-state reaction (conformational change) model, bivalent analyte and heterogeneous ligand model. The association and dissociation kinetic rate constants (k_a_, and k_d_, respectively) and the dissociation constant (K_D_), or equivalent, were estimated for each model.

### Mini-osmotic pump implantation

10-week-old, female C57Bl/6 (Crl) mice were obtained from Scanbur (Denmark) and randomly divided in 2 experimental groups; FnE (n=6) and saline control (n=6). Mice were housed in standard conditions with normal dark-light cycle. This method was approved by the ethical committee for animal experimentation in Uppsala (C45/13) and conform to the Animal Research: Reporting of In Vivo Experiments guidelines developed by the National Centre for the Replacement, Refinement and Reduction of Animals in Research. Animals were acclimated to their environment for 1 week and given *ad libitum* access to water and food. Mini-osmotic pumps (Alzet 1002, 0,25µl/hr) containing FnE (200ng/µl with 0,1% BSA in saline) or controls (0,1% BSA in saline) were implanted subcutaneously in mice anesthetized with isoflurane (Forene). Post-operative analgesia (Temgesic) was provided daily for 3 days. After exposure to 50 ng/hr FnE or saline (control) for 2 weeks, skin biopsies were collected around the pump using 4 mm biopsy punches. Tissues were fixed in 4% PFA and dehydrated and imbedded in paraffin before being sectioned in 5 um sections. Sections were de-parafinized and rehydrated before being blocked in 10% hydrogen peroxide and 1% BSA in PBS. Immunohistochemical staining for α-SMA (clone 1A4, Sigma) was performed using an HRP-coupled anti-mouse antibody with DAB as a substrate. Antibody incubation times were 1 h at room temperature. The samples were washed in TBS-T (0.1% Triton-X100) 3×10 minutes after primary and secondary antibody incubations. A standard hematoxylin and eosin staining was done to assess general morphology. Quantifications were performed blindly with ImageJ software by conversion to binary images after color de-convolution to separate DAB staining, as previously described (57).

### Statistics

Statistical analyses of cell cultured derived data was performed using two-sided one sample t-test, student’s t-test or analysis of variance (ANOVA) followed by Bonferroni-correction *post hoc* test, animal data was analyzed using the two-sided Mann-Whitney U-test. All statistical analyses were carried out in the GraphPad Prism 5 software (GraphPad software). Statistical significance was defined as p<0.05. Values are presented as mean±SEM, mouse data are additionally presented by individual dots.

### Study approval

The use of the mouse model and methods have been approved by the Uppsala Animal Experiment Ethics Board.

## Authorship details

P.F. Fuchs, F. Heindryckx, C. Calitz, F. Binet and S.M.Ø. Solbak have contributed to the design, performing experiments, interpretation of data and writing of the manuscript. H.U. Danielson has contributed to experiments, data interpretation and revising of the manuscript, J. Kreuger has contributed to concept and design, interpretation of data, writing and revising the intellectual content and P. Gerwins has contributed to concept and design, interpretation of data, writing and revising the intellectual content. All authors have contributed to revising and final approval of the submitted manuscript.

## Acknowledgements

This work was supported by grants from the Swedish Research Council (to P.G. and J.K.), the Swedish Cancer Society (to P.G., J.K. and F.H.), the Swedish Childhood Cancer Foundation (to P.G.), the Wenner-gren foundation (F.H.) and the Lions Cancer Research Foundation (to P.G.). P.F.F, has been supported in part by Olga Jönssons Foundation and Lennanders Foundation for the advancement of medical and scientific research.

